# Isopeptor: a tool for detecting intramolecular isopeptide bonds in protein structures

**DOI:** 10.1101/2024.12.24.630248

**Authors:** Francesco Costa, Rob Barringer, Ioannis Riziotis, Antonina Andreeva, Alex Bateman

## Abstract

**Motivation:** Intramolecular isopeptide bonds contribute to the structural stability of proteins, and have primarily been identified in domains of bacterial fibrillar adhesins and pili. At present, there is no systematic method available to detect them in newly determined molecular structures. This can result in mis-annotations and incorrect modelling.

**Results:** Here, we present Isopeptor, a computational tool designed to predict the presence of intramolecular isopeptide bonds in experimentally determined structures. Isopeptor utilizes structure-guided template matching via the Jess software, combined with a logistic regression classifier that incorporates Root Mean Square Deviation (RMSD) and relative solvent accessible area (rASA) as key features. The tool demonstrates a recall of 1.0 and a precision of 0.95 when tested on a Protein Data Bank (PDB) subset of domains known to contain intramolecular isopeptide bonds that have been deposited with incorrectly modelled geometries. Isopeptor’s python-based implementation supports integration into bioinformatics workflows, enabling early detection and prediction of isopeptide bonds during protein structure modelling.

**Availability and implementation:** Isopeptor is implemented in python and can be accessed via the command line, through a python API or via a Google Colaboratory implementation (https://colab.research.google.com/github/FranceCosta/Isopeptor_development/blob/main/notebooks/Isopeptide_finder.ipynb). Source code is hosted on GitHub (https://github.com/FranceCosta/isopeptor) and can be installed via the python package installation manager PIP.

## Introduction

Intramolecular isopeptide bonds typically form between lysine and asparagine/aspartate residues, catalyzed by a nearby aspartate or glutamate (Kang and Baker, 2011). When found in β-sandwich folds, they occur in CnaA-like domains (linking β-strands of opposing β-sheets) and CnaB-like domains (linking adjacent β-strands of the same β-sheet). These bonds stabilize bacterial surface proteins against various stresses, increasing their thermal, mechanical and proteolytic resistance (Kang & Baker, 2011). While biologically important, their detection has previously relied on bioinformatic analyses of CnaA-like and CnaB-like families, biophysical investigations and manual structure annotation during the building of models. In cases where researchers have built models without additional biophysical or bioinformatic context, intramolecular isopeptide bonds have not been modelled, leading to inaccurate depositions in the PDB (Figure.1). Isopeptor is a sequence-independent method that combines geometry-based template matching and machine learning to enable the automated identification of isopeptide bonds in large protein structure databases.

**Figure. 1:**
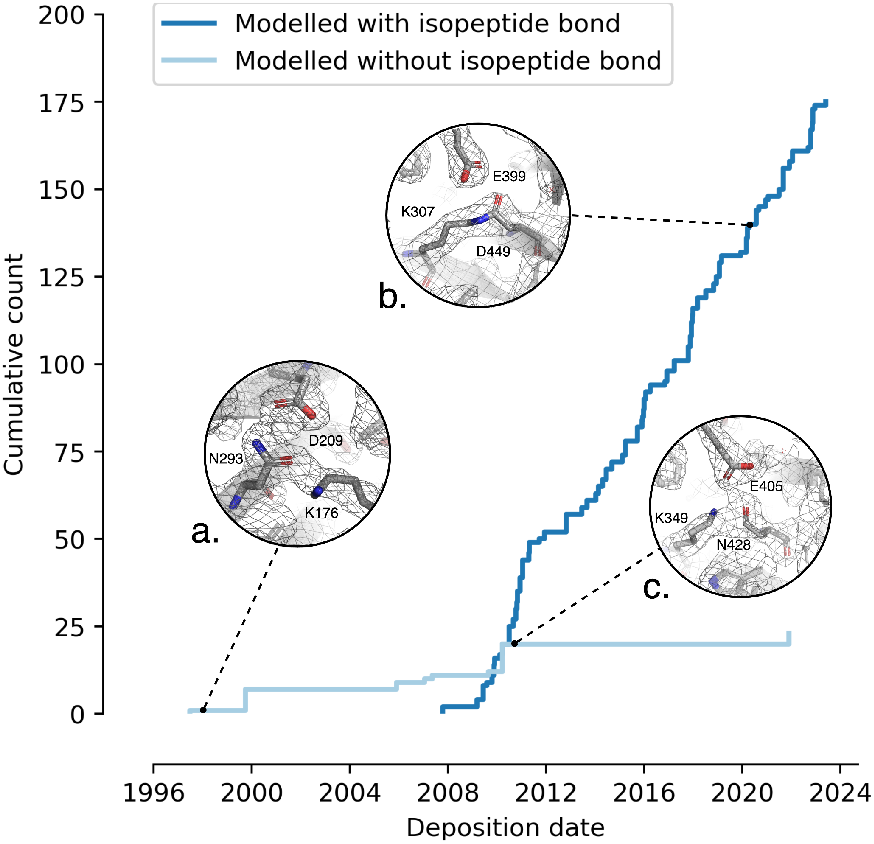
PDB depositions of intramolecular isopeptide bond domain structures over the years. Isopeptide bonds were not modelled into deposited structures until 2007, after Kang *et al*.’s discovery of intramolecular isopeptide bonds (Kang *et al*., 2007). A number of intramolecular isopeptide bond domain structures have been deposited without a modelled intramolecular isopeptide bond, or with bond geometries that do not fit the electron density map. A case of an unassigned isopeptide bond in a structure deposited prior to their discovery is shown (a. PDB ID: 1AMX) along with examples of correctly and incorrectly assigned isopeptide bond geometries (b. PDB ID:7K1U and c. PDB ID:2×9Y respectively), with their electron density maps (structural renderings generated with PyMol v3.1.1 https://pymol.org/).

**Figure. 2:**
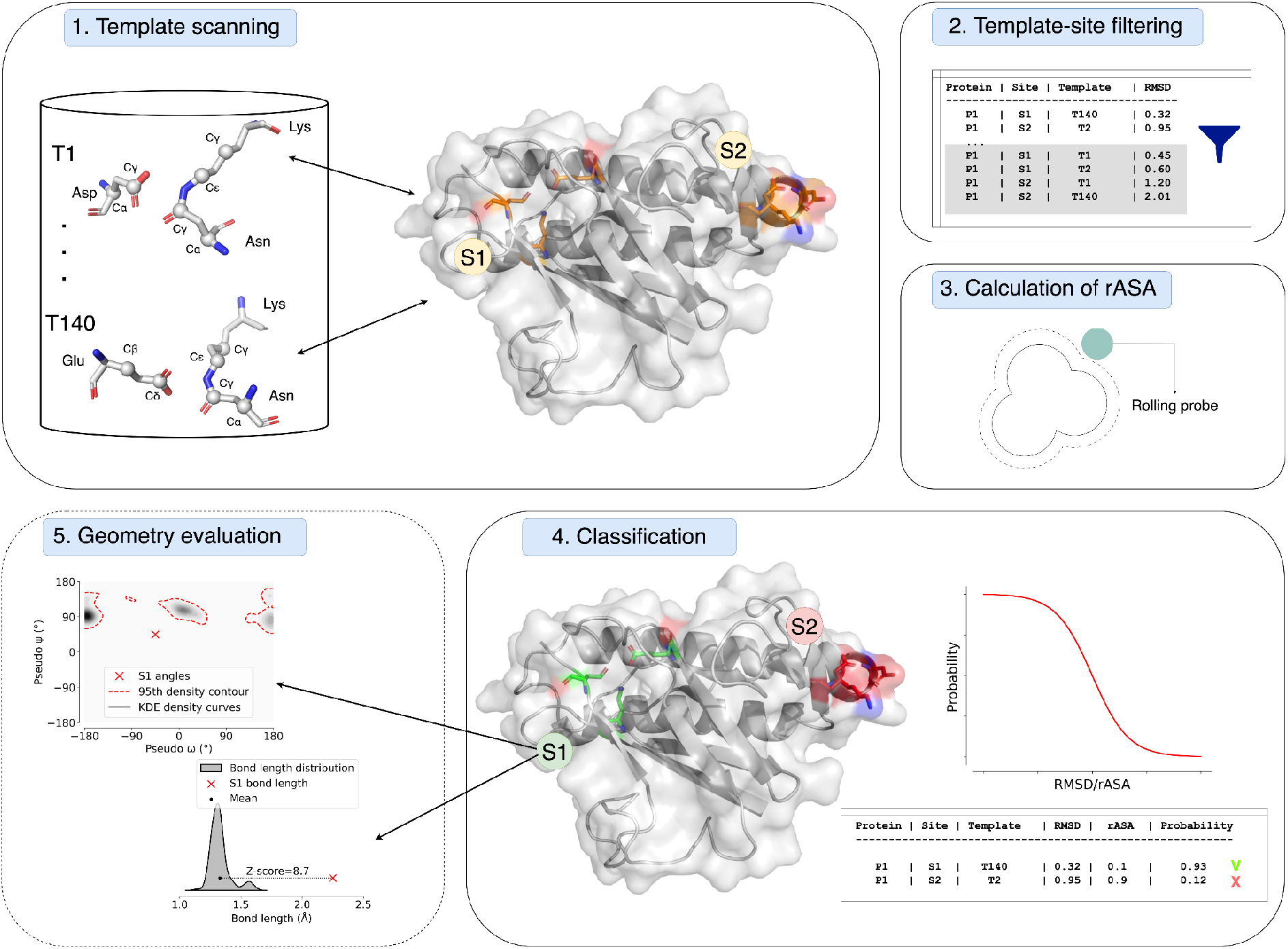
Isopeptor workflow. 1. Jess-based template scan is used to detect potential isopeptide bond signatures in target protein structures. A total of 140 templates coming from high quality structures are used for this purpose (labelled atoms are those included in the templates). Input structures can be in either PDB or CIF format. 2. If a site on the target protein structure matches with multiple templates, only the template-site match with the lowest RMSD is retained. 3. rASA is calculated. 4. Both RMSD with the closest template and rASA are taken as input by the logistic regression model for isopeptide bond classification which outputs a probability. 5. An optional final step is used to evaluate the geometry of isopeptide bonds. Two metrics are used to compare the parameters with the dataset of high quality structures: bond length Z-score and Kernel Density Estimate likelihood of dihedral angles. In the example shown, residues from site “S1” are predicted to form an intramolecular isopeptide bond but their geometry is sub-optimal. The output consists of a list of detected isopeptide bond signatures, each associated with an isopeptide bond probability and attributed to a topology (CnaA-like or CnaB-like) which is assigned depending on the topology of the closest template. The output can be stored in a tab-separated table. The image was generated with Drawio (https://app.diagrams.net/). Structure renderings were generated with PyMol version 3.1.1 (https://www.pymol.org/). The structure shown is PDB ID: 2×9Y, positions 447-625.

## Methods

### Dataset Creation

To develop Isopeptor, we compiled a curated training positive dataset consisting of 140 high confidence isopeptide bonds from the PDB through literature search and adopting a geometric template-based scan on the entire PDB (Barker and Thornton, 2003; Piehl and Burley, 2024). In some cases, we identified the presence of probable isopeptide bonds that were not correctly modelled in the deposited structure. For structures that were deposited with structure factor files and unmodelled intramolecular isopeptide bonds, structures were readjusted to fit an isopeptide bond to the electron density map using Molrep, Refmac and COOT (Vagin and Teplyakov, 1997; Murshudov *et al*., 2011; Emsley and Cowtan, 2004) within the CCP4i2 software suite (Potterton *et al*., 2018). Where necessary, we also defined the amino group of the isopeptide bond to belong to the N_Z_ of the lysine. We retained models demonstrating high resolution diffraction data (≤ 2.5Å) and excluded mutant structures lacking catalytic residues, or with catalytic residues demonstrating atypical geometries, such as the SpaC isopeptide bond from *Lactobacillus rhamnosus* (PDB ID: 6M48). For test purposes, we retained 19 isopeptide bond structures with incorrectly modelled geometries from high resolution PDB structures (i.e. uncorrected models with resolution ≤2.5 Å).

We also created a negative control dataset of 1,606 very high resolution (≤ 1.5Å) proteins of Eukaryotic origin from the PDB. We selected Eukaryote proteins because they do not (to our knowledge) present intramolecular isopeptide bonds, to minimise the risk of false negatives. Structures were chosen from 30% sequence identity PDB clusters (https://www.rcsb.org/) on 04/12/2024 to acquire a diverse range of proteins, excluding structures shorter than 80 amino acids. We split the negative dataset into two sets consisting of 75% of the initial dataset for training purposes and 25% of it to complement the positive test dataset.

### Template Matching

Jess is a program that enables structure-guided identification of target protein structures using customised templates consisting of three-dimensional atomic coordinates. We tested six template sets for the Lys-Asn-Asp, Lys-Asp-Glu, and Lys-Asn-Glu triads that characterise intramolecular isopeptide bonds (see Supplementary table.1).

The Jess output provides the Root Mean Square Deviation (RMSD) between templates and matching atoms. We used each set of templates to scan both negative and positive training datasets and considered only template matches with the lowest RMSD for each target group of residues. Additionally, we prohibited matches between templates and targets from the positive dataset sharing sequence identity above 30%. To achieve this, the pairwise sequence identity was obtained by aligning the sequences surrounding each isopeptide bond (20 amino acids upstream and downstream from the first and last residues of the triad forming the intramolecular isopeptide bond) with the *align_optimal* function from the python package Biotite (package version: 1.3.0, python version: 3.12.2) (Kunzmann and Hamacher, 2018). These settings have been adopted in all subsequent template scans.

We compared the performances of each template set by looking at the RMSD histogram overlaps between positive and negative distributions. We manually verified that entries from the negative set with a low RMSD did not contain any isopeptide bond by analysing their electron density maps. We found that templates from set III and set VI (supplementary table 1) are the best for discriminating between positives and negatives: with 0 and 2 entries overlapping between positive and negative RMSD distributions, respectively. We decided to adopt set VI templates because they contain less atoms, enabling a quicker computation.

### Feature Engineering and Classification

Isopeptor employs a logistic regression model to classify template matches. We trained the model on the training dataset using two features: 1. RMSD with the closest template (excluding template-matches with sequence identity above 30%) from set VI and 2. Relative Accessible Solvent Area (rASA) was incorporated into the model to assess how buried the matching residues are within the hydrophobic core, a known requirement for the formation of intramolecular isopeptide bonds (Kang and Baker, 2011). The rASA calculation was performed with the *sasa* function from the Biotite python package (version 0.39.0) with the *point_number* parameter set to 500. Rost and Sander maximum surface accessible values were used to calculate the rASA (Rost and Sander, 1994). The logistic regression model was developed using the *LogisticRegression* function from the Scikit-learn python package (version 1.3.0) with default parameters (https://scipy.org/).

### Implementation

Isopeptor is implemented in python and can be accessed via the command line, through a python API or via a Google Colaboratory implementation (https://colab.research.google.com/github/FranceCosta/Isopeptor_development/blob/main/notebooks/Isopeptide_finder.ipynb). Source code is hosted on GitHub (https://github.com/FranceCosta/isopeptor) and can be installed via the python package installation manager PIP.

## Results

### Validation

We used the entire set of intramolecular isopeptide bond-containing structures for the generation of the template dataset and subsequent training of the logistic regression classifier, as the positive dataset was not large enough for a training/test split (comprising only 28 clusters after 30% sequence identity clustering). For this reason, the performance of Isopeptor’s predictive model was evaluated on the test set of structures containing incorrectly modelled intramolecular isopeptide bonds (representing potential use cases where undetected isopeptide bonds are flagged by Isopeptor during protein structure modelling) and Eukaryotic depositions lacking intramolecular isopeptide bonds. Isopeptor was applied to this test set, correctly predicting all 19 isopeptide bonds with a probability above the decision threshold of 0.5, showing a recall of 1.0 and a precision of 0.95, indicating the presence of a low rate of false positives.

### Quality assessment metrics

We employed two metrics to assess the geometric quality of predicted isopeptide bonds: the bond length Z-score and the Kernel Density Estimations (KDE) likelihood of dihedral angles, calculated with the Scikit learn python package. The bond length Z-score represents the difference, expressed as the number of standard deviations, with the average isopeptide bond length. We define as outliers those values exceeding a Z-score of 4. We determined combinations of allowed dihedral angles by employing 6 different KDE models: one model for each of the three combinations of dihedral angles pairs (pseudo phi-pseudo psi, pseudo omega-pseudo psi, pseudo omega-pseudo phi) and the two isopeptide bond types (CnaA-like and CnaB-like). Given each KDE distribution, values outside the 95th percentile of the angles distribution from our dataset are defined as outliers.

## Discussion

Isopeptor represents a significant advance in the computational detection and validation of intramolecular isopeptide bonds in protein structures. By combining structure-guided template matching with machine learning classification, our tool achieves high precision and recall on experimentally determined structures with incorrectly modelled or un-modelled intramolecular isopeptide bonds. The integration of geometric quality assessment metrics provides valuable guidance for structure refinement, particularly in challenging cases involving low-resolution data. Its implementation as a python package enables the integration into existing structural biology workflows. We anticipate that Isopeptor will facilitate the identification and correct modeling of these important structural features, ultimately contributing to our understanding of protein stability mechanisms in bacterial surface proteins, and future protein folds that contain intramolecular isopeptide bonds. Future updates to the package will implement direct CIF input file parsing (currently handled by a CIF to PDB conversion step), to enable the parsing of very large protein structures.

## Author contributions

Francesco Costa (Conceptualization [equal], Data curation [equal], Formal analysis [equal], Software [lead], Visualization [lead], Writing—original draft [equal]), Rob Barringer (Conceptualization [equal], Data curation [equal], Formal analysis [lead], Writing—review & editing [equal]), Ioannis Riziotis (Methodology [equal], Conceptualization [equal], Writing—review & editing [equal]), Antonina Andreeva (Conceptualization [equal], Data curation [equal], Formal analysis [equal], Writing—original draft [equal]) and Alex Bateman (Conceptualization [equal], Funding acquisition [lead], Project administration [lead], Supervision [equal], Writing—original draft [equal], Writing—review & editing [equal])

## Funding

This work was supported by the UKRI Biotechnology and Biological Sciences Research Council [BB/X012492/1] and EMBL core funds, and by a UKRI Engineering and Physical Sciences Research Council Impact Accelerator Account (grant number EP/X525674/1).

## Data availability

The code to reproduce all data analysis performed in the publication is available at: https://github.com/FranceCosta/Isopeptor_development/. The dataset of intramolecular isopeptide bond-containing structures is available at: https://github.com/FranceCosta/Isopeptor_development/blob/main/data/tables/20241209_isopeptideBonds.csv while manually remodelled structures are found at: https://github.com/FranceCosta/Isopeptor_development/tree/main/data/PDB_bad_bonds_check_reassigned. All other data are available at: https://zenodo.org/records/14532222.

The method can be accessed via google colab: https://github.com/FranceCosta/Isopeptor_development/blob/main/notebooks/Isopeptide_finder.ipynb or run locally following the instructions provided at: https://github.com/FranceCosta/isopeptor.

## Notes

https://github.com/FranceCosta/isopeptor

https://zenodo.org/records/14532222

https://github.com/FranceCosta/Isopeptor_development/

### Competing Interest Statement

The authors have declared no competing interest.

## References

Barker, J.A. and Thornton, J.M. (2003) An algorithm for constraint-based structural template matching: application to 3D templates with statistical analysis. Bioinformatics, 19, 1644–1649.

Emsley, P. and Cowtan, K. (2004) Coot : model-building tools for molecular graphics. Acta Crystallogr D Biol Crystallogr, 60, 2126–2132.

Kang, H.J. et al. (2007) Stabilizing Isopeptide Bonds Revealed in Gram-Positive Bacterial Pilus Structure. Science, 318, 1625–1628.

Kang, H.J. and Baker, E.N. (2011) Intramolecular isopeptide bonds: protein crosslinks built for stress? Trends in Biochemical Sciences, 36, 229–237.

Kunzmann, P. and Hamacher, K. (2018) Biotite: a unifying open source computational biology framework in Python. BMC Bioinformatics, 19, 346.

Murshudov, G.N. et al. (2011) REFMAC 5 for the refinement of macromolecular crystal structures. Acta Crystallogr D Biol Crystallogr, 67, 355–367.

Piehl, D.W. and Burley, S.K. (2024) Parallel delivery of experimentally determined structures and computed structure models at RCSB protein data bank (RCSB PDB, RCSB.ORG). Biophysical Journal, 123, 280a.

Potterton, L. et al. (2018) CCP 4 i 2: the new graphical user interface to the CCP 4 program suite. Acta Crystallogr D Struct Biol, 74, 68–84.

Rost, B. and Sander, C. (1994) Conservation and prediction of solvent accessibility in protein families. Proteins, 20, 216–226.

Vagin, A. and Teplyakov, A. (1997) MOLREP : an Automated Program for Molecular Replacement. J Appl Crystallogr, 30, 1022–1025.

