## Supplementary information for "Isopeptor: a tool for detecting intramolecular isopeptide bonds in protein structures"

| Set | Bond residue 1 | Catalytic residue | Bond residue 2 | Number of template atoms | RMSD curves overlap |
|---|---|---|---|---|---|
| I | Lys$_{N\zeta}$ | Asn$_{O\delta1}$/Asp$_{O\delta1}$ | Glu$_{O\varepsilon1,O\varepsilon2}$/Asp$_{O\delta1,O\delta2}$ | 4 | 1063 |
| II | Lys$_{C\varepsilon,N\zeta}$ | Asn$_{O\delta1,C\gamma}$/Asp$_{O\delta1,C\gamma}$ | Glu$_{O\varepsilon1,O\varepsilon2,C\gamma}$/Asp$_{O\delta1,O\delta2,C\gamma}$ | 7 | 262 |
| III | Lys$_{C\gamma,C\delta,C\varepsilon}$ | Asn$_{C\alpha,C\beta,C\gamma}$/Asp$_{C\alpha,C\beta,C\gamma}$ | Glu$_{C\beta,C\gamma,C\delta}$/Asp$_{C\alpha,C\beta,C\gamma}$ | 9 | 0 |
| IV | Lys$_{C\delta,C\varepsilon}$ | Asn$_{C\beta,C\gamma}$/Asp$_{C\beta,C\gamma}$ | Glu$_{C\gamma,C\delta}$/Asp$_{C\beta,C\gamma}$ | 6 | 17 |
| V | Lys$_{C\gamma,C\delta}$ | Asn$_{C\alpha,C\beta}$/Asp$_{C\alpha,C\beta}$ | Glu$_{C\beta,C\gamma}$/Asp$_{C\alpha,C\beta}$ | 6 | 82 |
| VI | Lys$_{C\gamma,C\varepsilon}$ | Asn$_{C\alpha,C\gamma}$/Asp$_{C\alpha,C\gamma}$ | Glu$_{C\beta,C\delta}$/Asp$_{C\alpha,C\gamma}$ | 6 | 2 |

**Supplementary table.1**: Table showing tested combinations of atoms used in templates. Template set I consists of 4 atoms; the terminal nitrogen from the lysine (Lys$_{N\zeta}$), the terminal oxygen from the asparagine or aspartate (Asn$_{O\delta1}$/Asp$_{O\delta1}$), and the two terminal oxygens from either glutamate or aspartate (Glu$_{O\varepsilon1,O\varepsilon2}$/Asp$_{O\delta1,O\delta2}$). Template set II consists of 7 atoms; the terminal nitrogen and carbon from the lysine (Lys$_{C\varepsilon,N\zeta}$), the terminal oxygen and carbon from the asparagine or aspartate (Asn$_{O\delta1,C\gamma}$/Asp$_{O\delta1,C\gamma}$), and both terminal oxygens and terminal carbon from glutamate or aspartate (Glu$_{O\varepsilon1,O\varepsilon2,C\gamma}$/Asp$_{O\delta1,O\delta2,C\gamma}$). Templates from set III comprise 9 atoms in total: the three last carbons from lysine (Lys$_{C\gamma,C\delta,C\varepsilon}$), from asparagine or aspartate (Asn$_{C\alpha,C\beta,C\gamma}$/Asp$_{C\alpha,C\beta,C\gamma}$) and from glutamate or aspartate (Glu$_{C\beta,C\gamma,C\delta}$/Asp$_{C\alpha,C\beta,C\gamma}$). Sets IV and V are variations of set III, where respectively the first and last atoms for each set III residue atoms were excluded. Lastly, set VI is another variation of set III, where the central atom was excluded from each amino acid set. Templates were compared by looking at the number of overlapping hits between RMSD curves of positive and negative hits from the training set. Template-matches with sequence identity above 30% were excluded for this purpose.

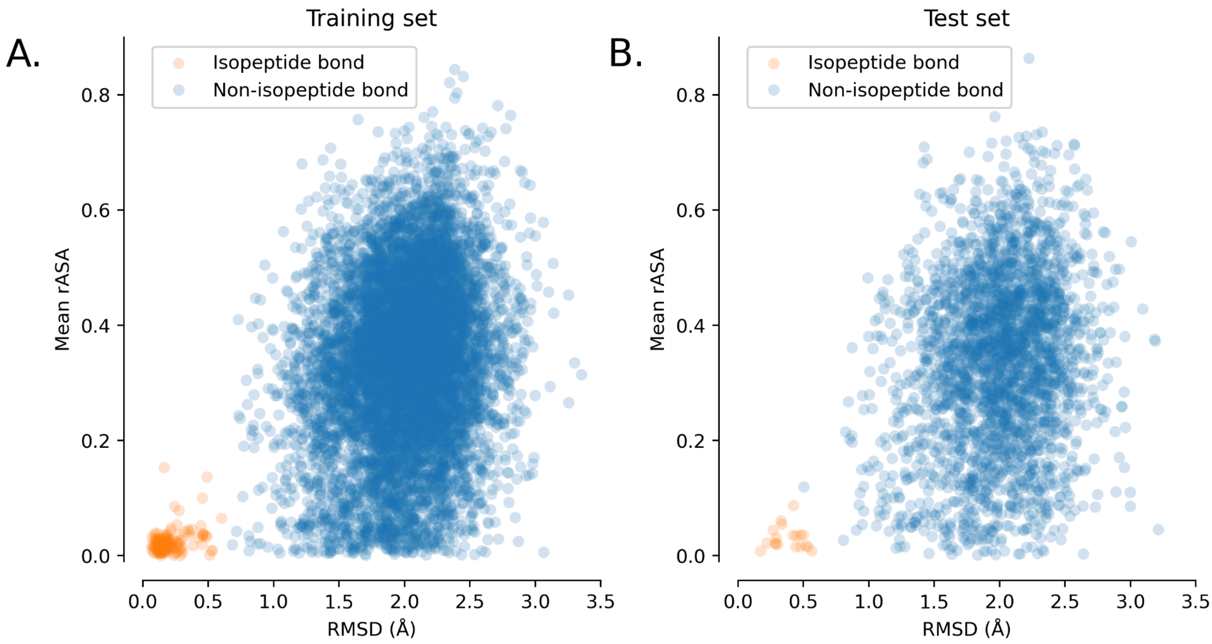


**Supplementary figure.S1**: Distribution of rASA average across match residues and RMSD (Å) with the closest template after the exclusion of template-matches with sequence identity above 30%. Data shown for training and test set (respectively A. and B.). The isopeptide bonds from the test set are all characterised by wrong or incorrect geometries which mostly affects terminal atoms from the lysine and asparagine/aspartate which form the isopeptide bond. This does not strongly influence the RMSD because it is not calculated considering terminal atoms (template set VI), therefore leading to very good performances of the logistic regression model.

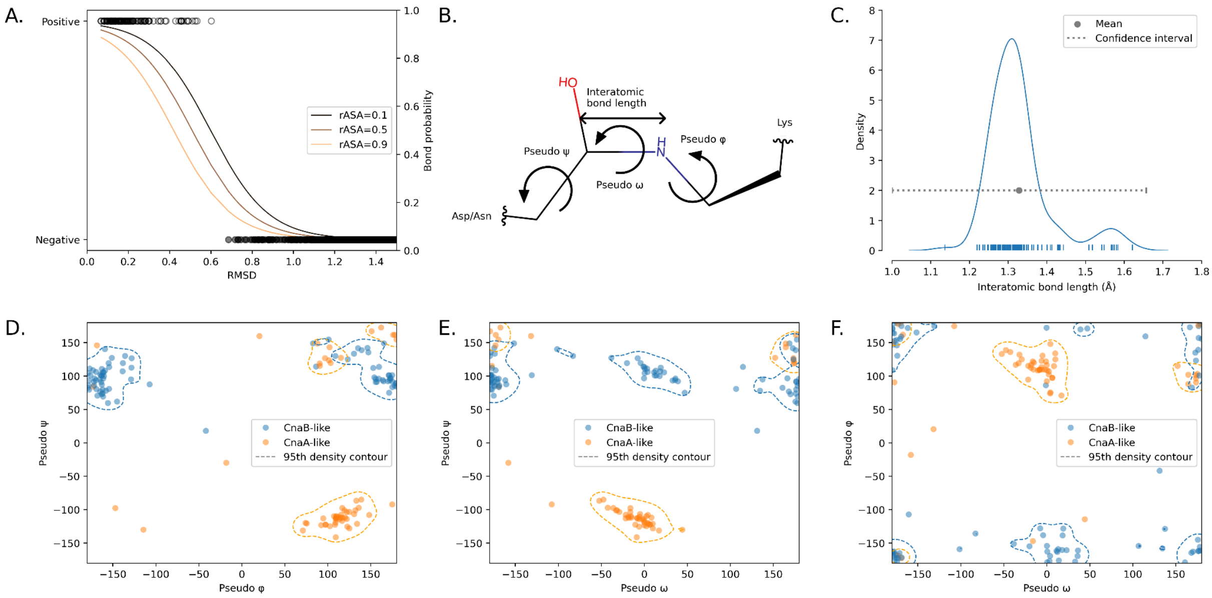


**Supplementary figure.S2**: A. Logistic regression probability curve as a function of RMSD and rASA. B. Isopeptide bond angles have been named after the peptide-bond dihedral angles nomenclature: pseudo φ is the angle around the bond between Asp/Asn$_{C\beta}$ and Asp/Asn$_{C\gamma}$, ω between Asp/Asn$_{C\gamma}$ and Lys$_{N\zeta}$ bond, and ψ between Lys$_{N\zeta}$ and Lys$_{C\varepsilon}$ bond. Image drawn with chemical-sketch (https://www.rcsb.org/chemical-sketch). C. Interatomic bond length of high-resolution structures containing intramolecular isopeptide bonds (≤2.5 Å). Values exceeding 4 standard deviations from the mean (confidence interval) have been excluded. The average interatomic isopeptide bond length is of 1.328 ± 0.082 Å, which is comparable with peptide bond distances (Allen *et al.*, 1987).
Distribution of pseudo φ-pseudo ψ ( D.), pseudo ω-pseudo ψ (E.) and pseudo ω-pseudo φ (F.) angles. Dotted lines represent the 95th percentile density contour calculated with a Kernel Density Estimate model for each combination of angle pairs and isopeptide bond type. Intramolecular isopeptide bonds' dihedral angles appear to differ from those tolerated by peptide bonds. Intramolecular bonds tolerate both *cis* and *trans* conformations, with CnaA-like domains preferring the trans conformation (62% of CnaA-like domains), and the CnaB-like domains preferring the cis conformation (60% of CnaB-like domains). This is in contrast to peptide bonds, which predominantly prefer a *trans* conformation (99.7% of peptide bonds, Weiss *et al.*, 1998). There are exceptions, for example in Xaa-Pro peptide bonds where *cis* and *trans* conformers are energetically equivalent, and *cis–trans* isomerization can significantly impact protein stability (Horng and Raines, 2006). It may therefore be the case that the preferred conformations of intramolecular isopeptide bonds within CnaA-like and CnaB-like domains represent energetically favourable conditions that increase domain stability. Analysis of pseudo- φ and ψ angles reveals significant differences between intramolecular isopeptide bonds and peptide bonds. CnaA-like *cis* bonds cluster in a region of the Ramachandran plot which is normally not populated in standard trans peptide bonds. Other regions usually characterised by Ramachandran outliers are conversely occupied by CnaB-like (both cis and trans) and by trans CnaA-like bonds.

### References

Allen,F.H. et al. (1987) Tables of bond lengths determined by X-ray and neutron diffraction. Part 1. Bond lengths in organic compounds. J. Chem. Soc., Perkin Trans. 2, S1.

Horng,J. and Raines,R.T. (2006) Stereoelectronic effects on polyproline conformation. Protein Science, 15, 74–83.

Weiss,M.S. et al. (1998) Peptide bonds revisited. Nat Struct Mol Biol, 5, 676–676.
